# Scutellarin inhibits ferroptosis by promoting cellular antioxidant capacity through regulating the Nrf2 signaling

**DOI:** 10.1101/2025.06.20.660655

**Authors:** Hai-yan Yang, On-kei Chan, Xiaodi Huang, Liang Yan, Nuo Sun, Ya-ping Li, Zi-jian Shi, Qing-bing Zha, Dong-yun Ouyang, Jinhua Li, Xian-hui He

## Abstract

Ferroptosis is a lytic form of regulated cell death that is driven by iron-dependent lipid peroxidation, and has been implicated in various diseases including acute kidney injury (AKI). Scutellarin is a flavonoid isolated from *Erigeron breviscapus* (Vant.) Hand.-Mazz. and possesses various pharmacological activities including anti-inflammatory and antioxidative properties. Yet it is unclear whether scutellarin can inhibit ferroptosis and mitigate related diseases. In this study, we found that scutellarin was able to inhibit ferroptosis in both human HK-2 cells and mouse bone marrow-derived macrophages stimulated with RSL3 or erastin. Mitochondrial dysfunction and reactive oxygen species generation were counteracted by scutellarin treatment, suggesting involvement of its antioxidative activity. Furthermore, scutellarin increased the nuclear levels of Nrf2 and the expression of its target genes including HO-1 and GPX4. Scutellarin-mediated inhibition of ferroptosis and an increase of these proteins was abrogated by the co-treatment with brusatol, an Nrf2 inhibitor, indicating an essential role for Nrf2 in this process. In a mouse model of folic acid-induced AKI, scutellarin mitigated acute renal damage as revealed by histopathological analysis and serum blood urea nitrogen and creatinine assays. Folic acid-induced acute renal injury was associated with increased ferroptosis as revealed by elevated levels of 4-hydroxynonenal (4-HNE), a surrogate marker of ferroptosis, which were diminished by scutellarin cotreatment. Specifically, the elevated 4-HNE levels in macrophages (MAC-2 positive) and other renal cells were suppressed by scutellarin. Collectively, scutellarin can inhibit ferroptosis both in cultured cells and in a mouse model of AKI by regulating Nrf2 signaling.

## Introduction

Ferroptosis is a nonapoptotic form of regulated cell death that is driven by iron-dependent lipid peroxidation [1–4]. The (phosphor)lipid peroxide-reducing enzyme glutathione peroxidase 4 (GPX4) is a central regulator of ferroptosis and prevents this cell death by catalyzing lipid peroxides into non-toxic lipid alcohols in the presence of sufficient glutathione [2, 5, 6]. The cystine/glutamate antiporter (X_c_^−^, (also known as xCT)) system is essential to maintain the levels of intracellular glutathione [6]. In addition to the system X_c_^−^/GPX4 cellular antioxidant axis, recent studies showed that ferroptosis suppressor protein 1 (FSP1) is the second mainstay of ferroptosis inhibiting antioxidant enzyme. Mechanistically, FSP1 is recruited to the plasma membrane by myristoylation and acts there as an oxidoreductase to reduce ubiquinone (also known as coenzyme Q_10_) into its reduced form ubiquinol, which functions as a lipophilic radical trapping antioxidant to block lipid peroxides [7, 8]. In parallel to mitochondrial GPX4, dihydroorotate dehydrogenase (DHODH) has been found to attenuate ferroptosis in mitochondria by reducing ubiquinone to ubiquinol [9]. Ferroptosis is also regulated by small antioxidative molecules such as α-tocopherol (vitamin E), which acts as lipophilic radical-trapping antioxidants to terminate the propagation of lipid peroxides [3, 5]. Moreover, two other recent reports showed that 7-hydrocholesterol is an endogenous suppressor of ferroptosis by acting as an antioxidant to protect against lipid peroxidation, thereby regulating the sensitivity of cells to ferroptosis [10, 11]. Notably, several important antioxidant proteins, such as GPX4 and heme oxygenase-1 (HO-1), are encoded by target genes of nuclear factor erythroid 2-related factor 2 (Nrf2), indicating a critical role for Nrf2 in modulating ferroptosis [12, 13]. Thus, the antioxidant systems of a cell have critical roles in protecting against ferroptosis.

As an oxidative form of cell death, ferroptosis can be induced by disruption of the cellular antioxidant systems through inhibition of GPX4 activity by genetic deletion of this gene or by RSL3-induced GPX4 inactivation [5]. This form of cell death can also be triggered by depletion of glutathione through inhibiting the system X_c_^−^ and thereby indirectly decreasing GPX4 activity [1, 5]. Although many studies adopted such biochemical methods to induce ferroptosis in vitro in cultured cells, emerging evidence implicates this oxidative form of regulated cell death in various pathological conditions including degenerative diseases, cancers, and organ injury [4, 6, 14]. Deletion of GPX4 results in acute renal failure in mice, indicating an essential role for the glutathione/GPX4 axis in suppressing lipid oxidation-induced acute kidney injury (AKI) and associated death [15, 16]. Several studies in mouse models of AKI further reveal a prominent role of ferroptosis in the pathology of renal tubular injury, showing that targeting ferroptosis could improve the survival of mice with AKI [16, 17]. Interestingly, 7-hydrocholesterol administration protected the kidney from ischemia-reperfusion injury through inhibiting ferroptosis [10]. Targeting ferroptosis is therefore a promising avenue for the treatment of renal injury during AKI.

Scutellarin is a flavonoid isolated from *Erigeron breviscapus* (Vant.) Hand.-Mazz. [18], a traditional medicinal herb that has long been used to treat paralysis after stroke and joint pain of rheumatoid arthritis by Yi minority people of southwestern China [19]. Scutellarin is the major active ingredient of this herbal medicine [18]. Breviscapine that containing ≥90% scutellarin has been clinically used for the treatment of cardiovascular and cerebrovascular diseases [20]. Other clinical studies showed that breviscapine can improve hypertension, hyperlipidemia, diabetic peripheral neuropathy, and diabetic nephropathy [19]. Such therapeutic properties of breviscapine have been attributed to the anti-inflammatory and antioxidative activities of scutellarin [18–20]. Interestingly, our previous studies showed that scutellarin not only inhibits canonical NLR family pyrin domain containing 3 (NLRP3) inflammasome activation but also suppresses noncanonical NLRP3 activation mediated by caspase-11, thus inhibiting pyroptosis [21, 22]. One recent study found that scutellarin inhibits pyroptosis by selective degradation of p30/gasdermin D through autophagy [23]. Furthermore, we recently found that scutellarin could inhibit PANoptosis in macrophages and thereby conferring protection against multi-organ injury in a mouse model of hemophagocytic lymphohistiocytosis that is associated with PANoptosis [24]. In light of the finding that ferroptosis is a prominent form of regulated cell death in AKI [16, 17], it is of interest to investigate whether scutellarin can affect ferroptosis both in vitro and in vivo in mouse models of AKI.

In this study, we found that the flavonoid scutellarin effectively inhibited ferroptosis in human and mouse cells in response to the suppression of GPX4 or the system X_c_^−^. This effect of scutellarin was mediated by upregulating the Nrf2 antioxidant capacity. Scutellarin was able to mitigate renal damage in a mouse model of AKI, which was associated with the suppression of ferroptotic signaling, likely by upregulating the Nrf2-GPX4 axis.

## Materials and methods

### Reagents and antibodies

Scutellarin (B21478) was purchased from Shanghai Yuanye Bio-Technology Co. (Shanghai, China), dissolved in dimethyl sulfoxide (DMSO) at a concentration of 100 mM, and stored at ‒80 °C. High-glucose Dulbecco’s Modified Eagle Medium (DMEM) (C11995500BT), DMEM/F12 (C11330500BT), fetal bovine serum (FBS) (10099141C), penicillin-streptomycin (15140122), C11-BODIPY 581/591 (D3861), and MitoSOX Red (M36008) were obtained from Thermo Fisher (Carlsbad, CA, USA). Folic acid (F7876), DMSO (D8418), propidium iodide (PI) (P4170), Hoechst 33342 (B2261), Tween-80 (P8074), Tween-20 (P1379), DL-dithiothreitol (DTT) (D0632) and CF488-conjugated goat-anti-mouse IgG (SAB4600237) were purchased from Sigma-Aldrich (St. Louis, MO, USA). Nrf2 (#12721), lamin A/C (#4777), β-tubulin (2128), β-actin (#3700) and HO-1 (#43966) were obtained from Cell Signaling Technology (Danvers, MA, USA). The antibody against GPX4 (ab125066) was bought from Abcam (Cambridge, UK). Anti-4-hydroxynonenal (4-HNE) antibody (JAI-MHN-020P) was purchased from AdipoGen (Liestal, Switzerland). AlexaFluor647 conjugated anti-mouse/human MAC-2 antibody was obtained from BioLegend (San Diego, CA, USA). Erastin (S7242), brusatol (S7956) and RSL3 (S8155) were purchased from Selleck (Houston, TX, USA). Ferrostatin-1 (HY-100579) was obtained from MedChemExpress (New Jersey, USA). WST-1 reagent (11644807001) was purchased from Roche (Mannheim, Germany). Nuclear and cytoplasmic protein extraction kit (P0027) was obtained from Beyotime (Sanghai, China). Creatinine assay kit (C011-2-1) was purchased from Nanjing Jiancheng Bioengineering (Nanjing, China). Blood urea nitrogen (BUN) kit (E2020) was obtained from Applygen (Beijing, China).

### Animals

C57BL/6J mice (6-8 weeks of age) were obtained from Guangzhou Ruige Biological Technology (Guangzhou, China). All mice were housed under controlled conditions at 24 ± 2 °C with a 12/12 h light-dark cycle, provided with ad libitum access to food and water. The animal experimental procedures were approved by the Jinan University Laboratory Animal Welfare and Ethics Committee and conducted in accordance with the committee’s guidelines.

### Cell culture and treatments

Human renal proximal tubular epithelial cell line HK-2 was a gift from Dr. Liang Yan (the First Affiliated Hospital of Jinan University, Guangzhou, China). Cells were maintained in DMEM/F12 supplemented with 10% FBS and 1% penicillin-streptomycin (complete DMEM/F12), with subculturing at a 1:3 or 1:4 ratio every 2-3 days. For experiments, HK-2 cells were seeded at 1.5×10^5^ cells/well (0.5 mL) in 24-well plates or 6×10^5^ cells/well (1.7 mL) in 6-well plates using complete DMEM/F12 medium and incubated overnight at 37 °C.

Bone marrow-derived macrophages (BMDMs) were isolated and cultured as previously described [25]. Briefly, bone marrow cells were flushed from femurs and tibias with 20 mL sterile PBS, centrifuged at 1500 rpm for 5 min at 4 °C, and resuspended in BM-Mac medium (80% complete DMEM + 20% M-CSF-conditioned medium from L929 cells). Cells were cultured in 10 cm dishes with 10 mL of BM-Mac medium for 6 days in a humidified 37 °C incubator with 5% CO₂, supplemented with 5 mL fresh BM-Mac medium on day 3. For experiments, BMDMs were seeded at 2.5×10^5^ cells/well (0.5 mL) in 24-well plates or 1.6×10^6^ cells/well (1.7 mL) in 6-well plates using complete DMEM and incubated overnight at 37 °C.

### Cytotoxicity assay

The cytotoxicity of scutellarin was evaluated by using WST-1 assay. In brief, BMDMs (2.5 × 10^5^/well) or HK-2 cells (3 × 10^4^/well) were seeded in 96-well plates (100 μL/well) and cultured at 37 °C under 5 % CO₂ overnight. The cells were then treated with different concentrations of scutellarin for 24 h. Thereafter, 10 μL of WST-1 was added to each well and incubated at 37 °C for 2 h. The absorbance at 450 nm was measured, and cell viability was evaluated and presented as percentages of control.

### Cell death assay

Cell death was determined as described previously [26]. BMDMs or HK-2 cells were seeded in 24-well plates overnight. After indicated treatments, the cells were incubated with freshly prepared staining solution containing PI (2 μg/mL) and Hoechst 33342 (5 μg/mL) for 10 min (0.5 mL/well, 37 °C, 5% CO₂). Fluorescence imaging was performed using an inverted fluorescence microscope with Rhodamine (PI signal) and DAPI (Hoechst signal) channels. Images of 5 random fields were captured for analysis. Percentages of cell death were determined by calculating the ratio of PI-positive nuclei (indicating dead cells) to Hoechst 33342-positive nuclei (total cells).

### Western blot analysis

Western blotting was performed as described previously [24]. Proteins from whole-cell lysates or nuclear proteins isolated by using the nuclear protein extraction kit (P0027, Beyotime) were separated by SDS-PAGE and transferred onto PVDF membranes (03010040001; Roche, Mannheim, Germany). Membranes were blocked with blocking buffer for 1 h at room temperature, followed by incubation with primary antibodies: Nrf2 (1:1000), HO-1 (1:2000), and GPX4 (1:8000). β-Actin (1:2000) was detected as a loading control. After overnight incubation at 4 °C, the membranes were washed with PBS containing 0.05% Tween-20 (PBST) and incubated with an HRP-conjugated secondary antibody for 1 h at room temperature. Protein bands were visualized using an enhanced chemiluminescence kit (BeyoECL Plus, P0018; Beyotime, Shanghai, China) and X-ray film (Carestream, Xiamen, China). Imaging and densitometric analysis were performed using a FluorChem 8000 system (AlphaInnotech, San Leandro, CA, USA) with ImageJ and GraphPad Prism 7.0 software (GraphPad Software Inc; San Diego, CA, USA).

### Measurement of mitochondrial superoxide

Mitochondrial superoxide levels (reflected by mitochondrial ROS (mtROS)) were assessed by staining with 3 μM of MitoSOX Red in live cells. After 15 min incubation, fluorescence signals were visualized using an Axio Observer D1 fluorescence microscope (Carl Zeiss MicroImaging GmbH, Gottingen, Germany).

### Assessment of mitochondrial membrane potential

Mitochondrial membrane potential (MMP) was evaluated by using 5 μg/mL of JC-1 (C2006; Beyotime) or TMRE kit (C2001S; Beyotime) according to the protocols of the supplier. Cells were incubated with the fluorophores for 30 min, respectively, followed by fluorescence imaging using an Axio Observer D1 microscope (Carl Zeiss).

### Lipid peroxidation product determination

Lipid peroxidation, a hallmark of ferroptosis, was assessed by staining with 2 μg/mL of C11-BODIPY 581/591. Cells were incubated with the fluorescent probe for 30 min, followed by fluorescence imaging using an Axio Observer D1 microscope (Carl Zeiss).

### Animal model of AKI

AKI was induced in female C57BL/6J mice through folic acid (FA) administration. The mice were randomly divided into four groups (*n* = 5 per group): vehicle control, scutellarin alone, FA alone, and scutellarin + FA combination. Scutellarin was prepared in PBS containing 2% Tween-80 as vehicle. The scutellarin and scutellarin + FA groups received oral gavage of scutellarin solution (200 mg/kg body weight) once a day for two consecutive days. On day 3, these groups received an additional scutellarin dose 3 h prior to intraperitoneal FA injection (250 mg/kg body weight). The FA and scutellarin + FA groups were then administered with FA solution dissolved in 0.3 M of NaHCO_3_ solution. Twenty-four hours post-FA injection, mice were anesthetized with ethyl ether for orbital blood collection. Blood samples were used to prepare serum by centrifugation (1300 rpm, 4 °C, 30 min) after incubation at room temperature for 60 min followed by overnight incubation at 4 °C. After euthanasia via cervical dislocation, kidney tissues were harvested for subsequent analysis.

### Serum creatinine and BUN measurement

Serum creatinine and BUN levels were determined using commercial assay kits according to the manufacturers’ protocols.

### Immunofluorescence staining

Left kidneys were fixed in 4% paraformaldehyde for 24 h (fully submerged) and were frozen sectioned. Tissue sections were stored at ‒80 °C until use. For antigen retrieval, sections were equilibrated in PBS (10 min, room temperature) followed by immersion in preheated sodium citrate buffer (10 mmol/L, pH 6.0, 80 °C, 30 min). After cooling to room temperature, sections underwent three PBS washes (5 min each) and were circumscribed with an immunohistochemical pen. Blocking was performed using PBS containing 5% goat serum and 0.1% Triton X-100 (60 min, room temperature). Primary antibodies were applied within demarcated areas and incubated at 4 °C overnight. Following PBS washes, corresponding fluorescent secondary antibodies were added (1 h, room temperature). After Hoechst 33342 nuclear staining (5 μg/mL PBS, 15 min), slides were mounted with anti-fade medium. Images were captured using a Zeiss AxioCam MR R3 CCD camera controlled by ZEN software (Carl Zeiss).

### Histochemical analysis

Right kidneys were fixed in 10% neutral formalin for 24 h (fully submerged) and paraffin sections were routinely prepared. Images of tissue sections were captured by using an Axio Observer D1 microscope (Carl Zeiss).

### Statistical analysis

Experiments were performed three times independently. Data are presented as mean ± standard deviation (SD). Statistical significance was analyzed using GraphPad Prism 7.0 (GraphPad Software Inc, San Diego, CA, USA). One-way ANOVA with Bonferroni *post hoc* test and unpaired Student’s *t*-test were applied for multi-group and two-group comparisons, respectively. Significance thresholds were defined as \**P* < 0.05, \*\**P* < 0.01, and \*\*\**P* < 0.001.

## Results

### Scutellarin inhibits ferroptosis both in mouse and human cells

As the antioxidant GPX4 and X_c_^−^ system have essential roles in regulating ferroptosis, the pharmacological inhibition of GPX4 and X_c_^−^ by RSL3 and erastin respectively can induce this form of cell death [1, 5]. Thus, we assessed the effects of scutellarin on ferroptosis in human renal proximal tubule epithelial cell line HK-2 and mouse BMDMs treated with RSL3 or erastin. PI staining was used to detect cell death in these cells. RSL3 or erastin treatment markedly increased the ratios of PI-positive cells (indicative of dying cells), whereas scutellarin co-treatment significantly decreased the levels of PI-positive cells (**Figure 1A-D**). RSL3- or erastin-induced cell death was blocked by co-treatment with ferrostatin 1 (Fer-1), featuring ferroptotic cell death. Similar results were obtained from BMDMs (**Figure 2A-D**). Scutellarin (200 μM) alone did not induce cell death in both HK-2 cells and BMDMs, which was verified by cytotoxicity assay showing that scutellarin was non-toxic at 200 μM or less (supplementary **Figure S1A, B**). These results together indicate that scutellarin is able to inhibit ferroptosis in both human and mouse cells.

**Figure 1.**
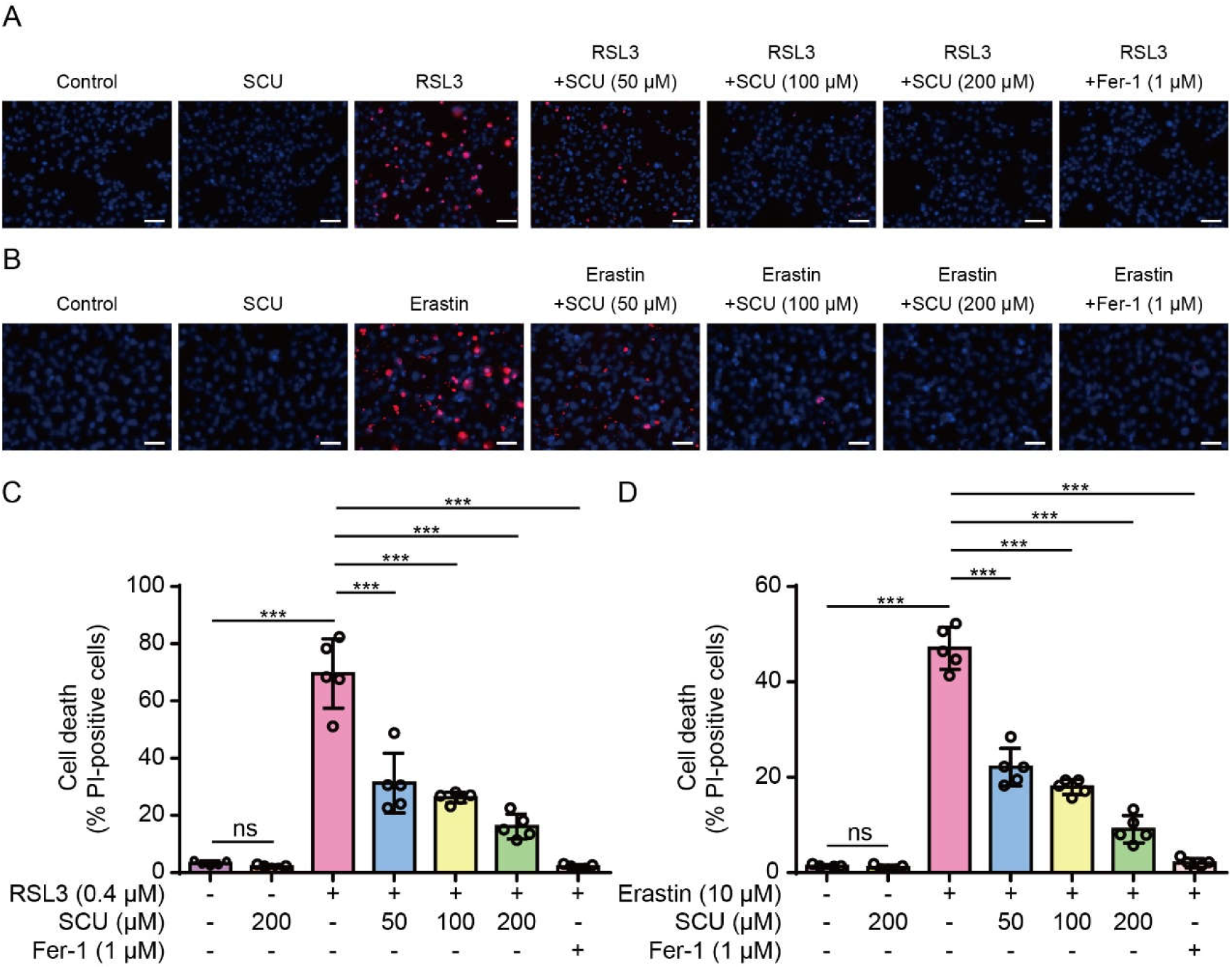
RSL3- or erastin-induced ferroptosis in HK-2 cells by is inhibited by scutellarin. HK-2 cells were pretreated with or without graded concentrations of scutellarin (SCU) for 1 h, followed by treatment with RSL3 (0.4 µM) for 5 h (A, C) or with erastin (10 µM) for 5 h (B, D). Cell death was assessed by propidium iodide (PI) staining (red, indicating dying cells) and Hoechst 33342 (blue, staining all nuclei). (A, B) Representative images captured by using a fluorescence microscope. Scale bars, 50 µm. (C, D) Quantitative analysis of percentages of cell death. PI-positive cells were quantified in five random fields, and the percentage of cell death was presented as the ratio of PI-positive cells to total cells (revealed by Hoechst 33342 staining). Data are presented as mean ± standard deviation (SD) (*n* = 5). \*\*\**P* < 0.001; ns, not significant.

**Figure 2.**
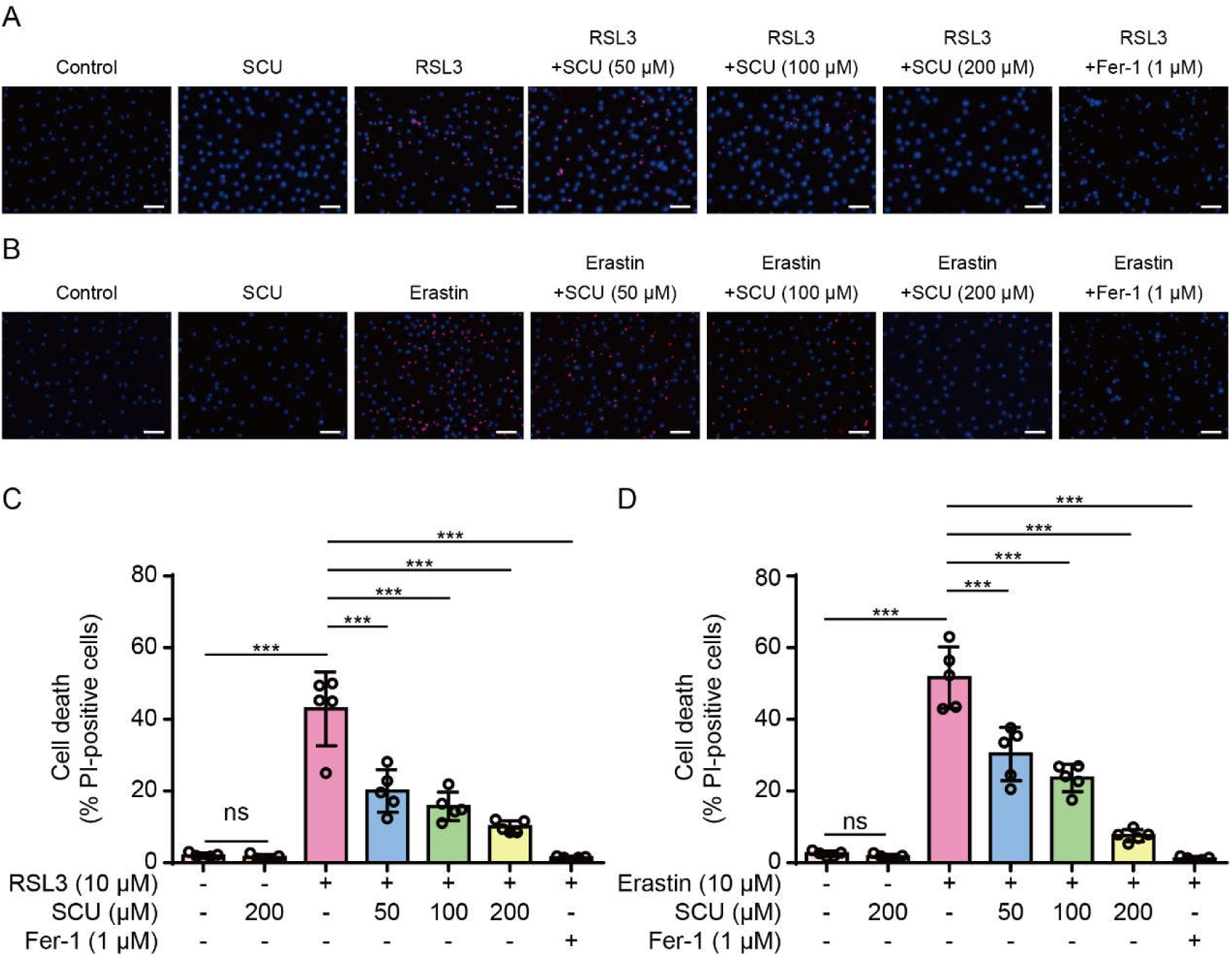
Ferroptosis in macrophages induced by RSL3 or erastin is inhibited by scutellarin. Bone-marrow-derived macrophages (BMDMs) were pretreated with or without graded concentrations of SCU for 1 h, followed by treatment with RSL3 (10 µM) for 24 h (A, C) or with erastin (10 µM) for 24 h (B, D). Cell death was assessed by propidium iodide (PI) staining (red, indicating dying cells) and Hoechst 33342 (blue, staining all nuclei). (A, B) Representative images captured by using a fluorescence microscope. Scale bars, 50 µm. (C, D) Quantitative analysis of percentages of cell death. PI-positive cells were quantified in five random fields, and the percentage of cell death was presented as the ratio of PI-positive cells to total cells (revealed by Hoechst 33342 staining). Data are presented as mean ± SD (*n* = 5). \*\*\**P* < 0.001; ns, not significant.

We next assessed the effect of scutellarin on lipid peroxidation, which is a hallmark of ferroptosis [2, 3]. C11-BODIPY 581/591 dye is a sensor for lipid peroxidation [27]: C11 can excited with 581 nm to emit red fluorescence (591 nm) at its reduced form; upon oxidation, the emission of red fluorescence is diminished whereas the green fluorescence (510 nm) excited by 488 nm is increased [28]. Fluorescence microscopy showed that RSL3 markedly increased green fluorescence and decreased red fluorescence in HK-2 cells and that these changes were reversed by co-treatment with scutellarin (**Figure 3A, C, E**). Similar results were observed in BMDMs (**Figure 3B, D, F**). Together, these results indicate that scutellarin can inhibit lipid peroxidation and ferroptosis in human and mouse cells.

**Figure 3.**
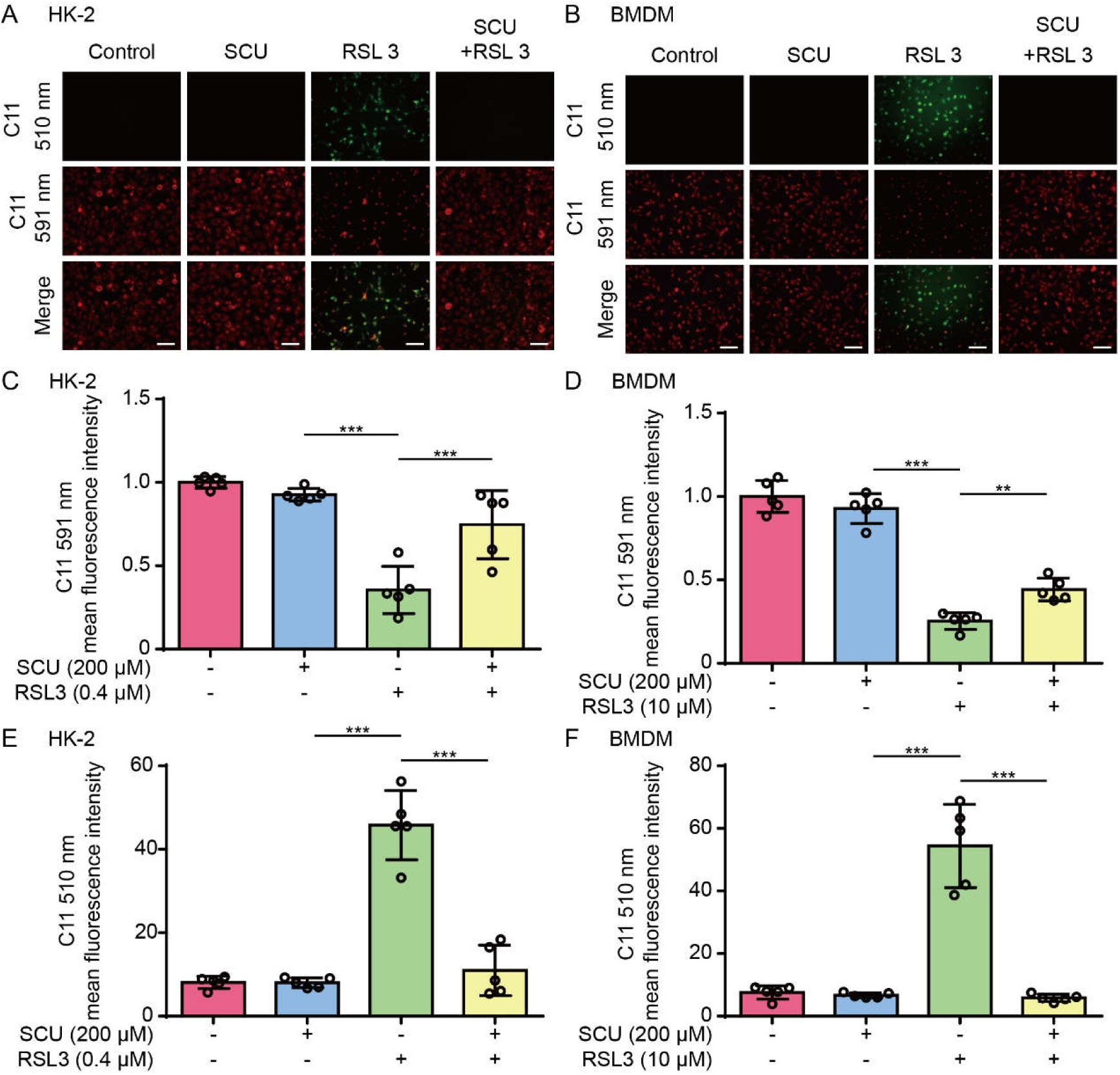
Lipid peroxidation in HK-2 and macrophages induced by RSL3 is inhibited by scutellarin. HK-2 cells or BMDMs were pretreated with or without SCU (200 μM) for 1 h, followed by treatment with RSL3 (0.4 µM) for 5 h (A, C, E) or with RSL3 (10 µM) for 24 h (B, D, F), respectively. Lipid peroxidation was assessed by using C11-BODIPY581/591 (5 µM) staining. Cells labeled with C11-BODIPY581/591 were excited at 488 nm and 561 nm to observe green and red fluorescence, respectively. (A, B) Representative images captured with a fluorescence microscopy. Scale bars, 50 µm. (C, D) Quantitative analysis of red fluorescence intensity of (A, B). (E, F) Quantitative analysis of green fluorescence intensity of (A, B). Data are presented as mean ± SD (*n* = 5). \*\**P* < 0.01; \*\*\**P* < 0.001.

### Scutellarin decreases mitochondrial damage during induction of ferroptosis

In view of the findings that mtROS have a crucial role in lipid peroxidation leading to ferroptosis [2, 9, 29–31], we next explored the effect of scutellarin on mtROS and mitochondrial function. Fluorescence microscopy showed that TMRE fluorescence (indicative of MMP) was decreased in HK-2 cells upon RSL3 treatment while co-treatment with scutellarin reversed this decrease (**Figure 4A, B**). JC-1 staining further confirmed a decrease of MMP in HK-2 cells upon RSL3 stimulation as revealed by decreased red fluorescence (indicating aggregated JC-1 within mitochondria) and increased green fluorescence (indicative of diffused JC-1 monomer in the cytosol), which was counteracted by scutellarin (**Figure 4C, D**). We also examined the production of mtROS by using MitoSOX staining and found that scutellarin significantly decreased MitoSOX fluorescence in HK-2 cells in response to RSL3 stimulation (**Figure 4E, F**). Similar results were obtained from BMDMs (**Figure 5A-F**). These results together indicate that scutellarin protects mitochondrial function and inhibits mtROS production during induction of ferroptosis.

**Figure 4.**
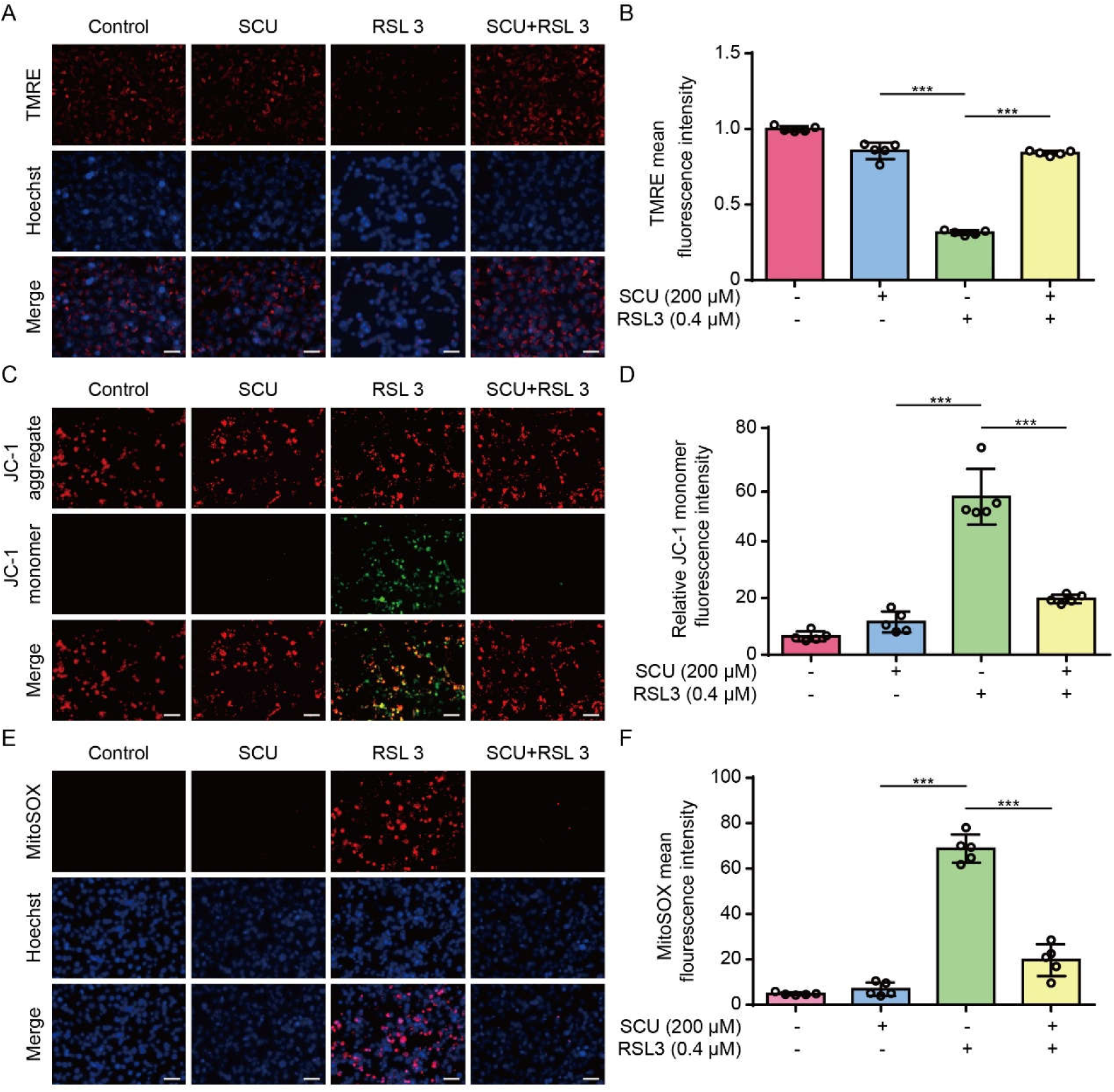
Mitochondrial dysfunction in HK-2 cells induced by RSL3 is inhibited by scutellarin. HK-2 cells were pretreated with different concentrations of SCU for 1 h, followed by treatment with RSL3 (0.4 µM) for 5 h. Mitochondrial membrane potential (MMP) was analyzed using TMRE (A, B) or JC-1 (C, D) staining. (A, B) Representative fluorescence images. (C, D) Quantitative analysis of TMRE and JC-1 in (A, B). (E, F) Mitochondrial superoxide (mtROS) was measured by MitoSOX staining and images were obtained using a fluorescence microscope (E). (F) Quantitative analysis of MitoSOX fluorescence. Data are presented as mean ± SD (*n* = 5). Scale bars, 50 µm. \*\*\**P* < 0.001.

**Figure 5.**
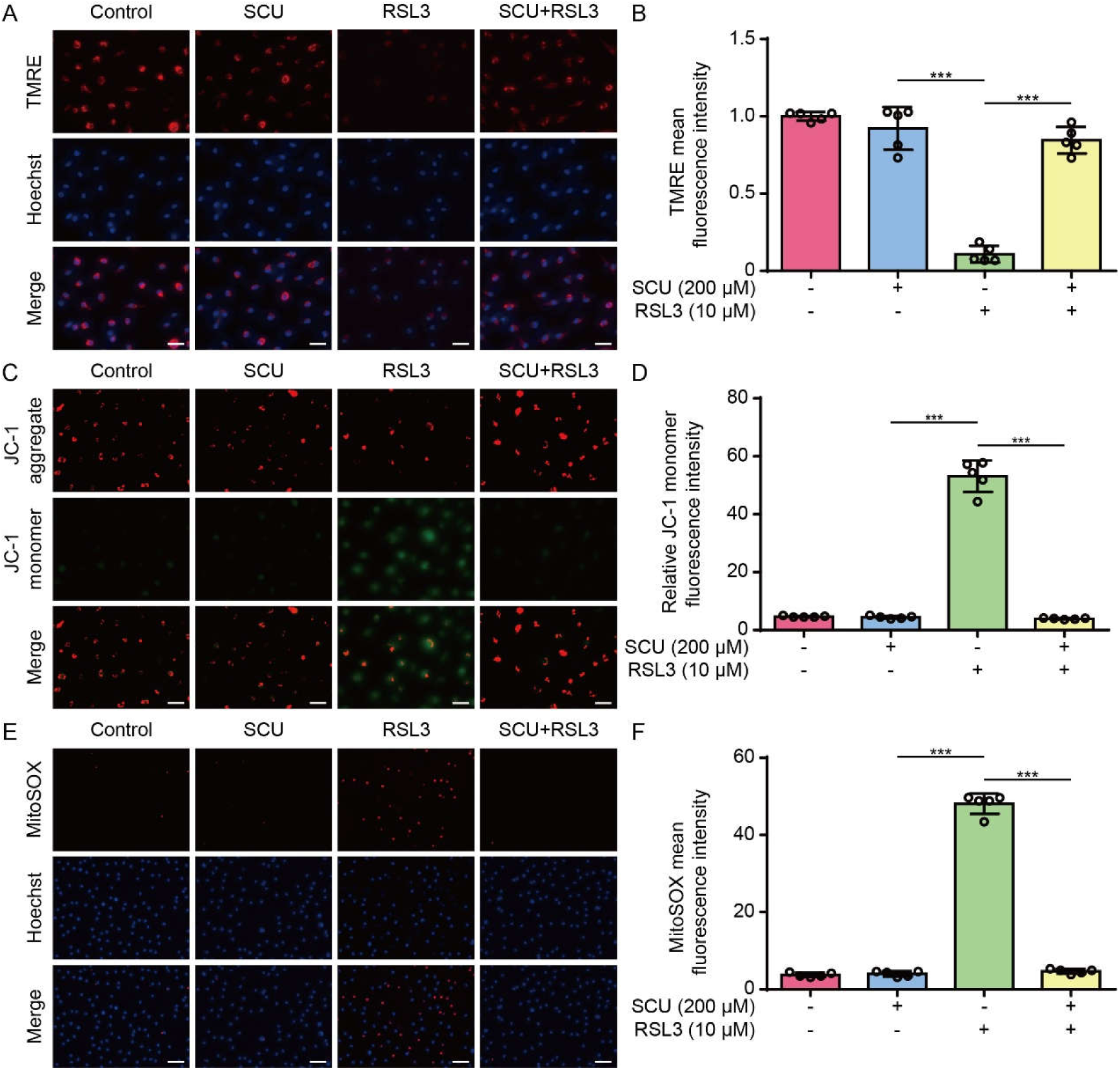
Mitochondrial dysfunction in macrophages induced by RSL3 is inhibited by scutellarin. BMDMs were pretreated with different concentrations of SCU for 1 h, followed by treatment with RSL3 (10 µM) for 24 h. Mitochondrial membrane potential (MMP) was analyzed using TMRE (A, B) or JC-1 (C, D) staining. (A, B) Representative fluorescence images. (C, D) Quantitative analysis of TMRE and JC-1 in (A, B). (E, F) Mitochondrial superoxide (mtROS) was measured by MitoSOX staining and images were obtained using a fluorescence microscope (E). (F) Quantitative analysis of MitoSOX fluorescence. Data are presented as mean ± SD (*n* = 5). Scale bars, 50 µm. \*\*\**P* < 0.001.

### Inhibition of ferroptosis by scutellarin relies on the upregulation of Nrf2 signaling

As the Nrf2 transcription factor is known to control the expression of many antioxidative proteins including HO-1 and GPX4 [12, 13], we next sought to investigate whether scutellarin could affect the expression of GPX4 through modulating Nrf2 activity. Accompanying inhibition of ferroptotic cell death (**Figure 1**), Western blotting showed that scutellarin counteracted the decrease of GPX4 induced by either RSL3 or erastin in HK-2 cells (**Figure 6A-D**). Consistently, scutellarin markedly increased the levels of Nrf2 expression as well as the HO-1 in HK-2 cells treated with RSL3 or erastin (**Figure 6A-D**). Similar results were derived from BMDMs (**Figure 6E-H**). We further assessed whether scutellarin could affect nuclear translocation of Nrf2 by detecting its levels in isolated nuclear fraction. Western blotting showed that scutellarin co-treatment with RSL3 or erastin dose-dependently increased Nrf2 levels in the nuclear fraction of HK-2 cells and BMDMs (supplementary **Figure S2A-H**). These results suggested that scutellarin increased the expression of GPX4 by upregulating the Nrf2 signaling.

**Figure 6.**
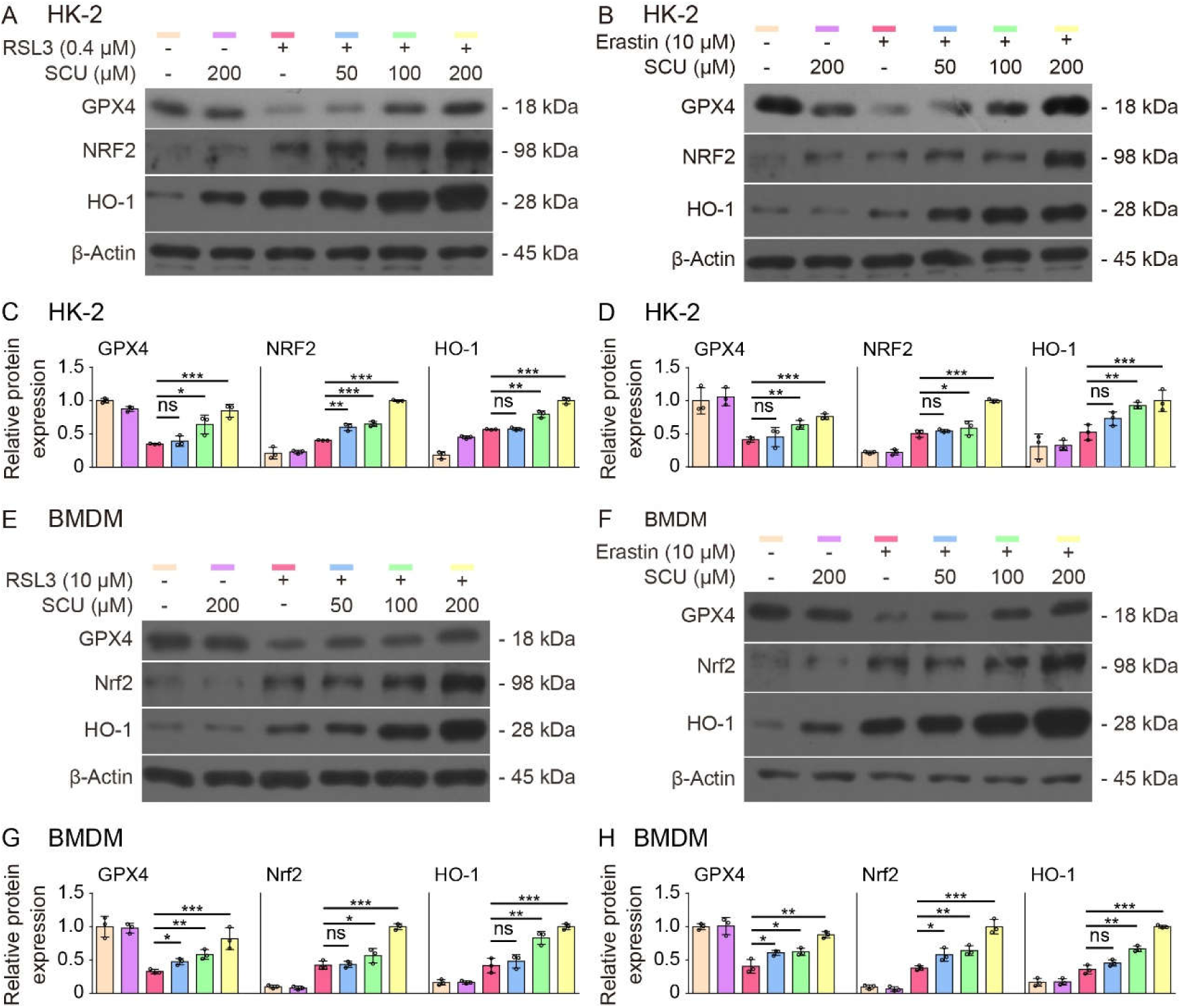
Scutellarin-mediated inhibition of ferroptosis is associated with upregulation of Nrf2 signaling. Cells were pretreated with different concentrations of SCU for 1 h, followed by treatment with RSL3 or erastin for 5 h (for HK-2) or for 24 h (for BMDMs). (A, B, E, F) Western blot analysis of Nrf2, HO-1 and GPX4 levels in cell lysates. β-Actin was used as a loading control. (C, D, G, H) Quantitative analysis of protein expression in (A), (B), (E), (F), respectively. Data are presented as mean ± SD (*n* = 3). \**P* < 0.05; \*\**P* < 0.01; \*\*\**P* < 0.001; ns, not significant.

To further verifying the role of Nrf2 in mediating the inhibitory effect of scutellarin on ferroptosis, we co-treated scutellarin with brusatol, an Nrf2 inhibitor that enhanced Nrf2 degradation [32], in HK-2 cells during induction of ferroptosis. PI staining showed that brusatol did not increased RSL3-induced cell death but significantly reversed scutellarin-mediated inhibition of RSL3-induced ferroptotic cell death (**Figure 7A, B**). Consistent with this, scutellarin-mediated increase of NRF2, GPX4 and HO-1 in RSL3-treated cells was counteracted by brusatol co-treatment (**Figure 7C, D**). We further assessed NRF2 levels in the nuclear fraction and found that scutellarin-induced NRF2 nuclear translocation was markedly suppressed by brusatol (**Figure 7E, F**). These results indicate that scutellarin inhibits ferroptosis in HK-2 cells by upregulating the Nrf2 signaling.

**Figure 7.**
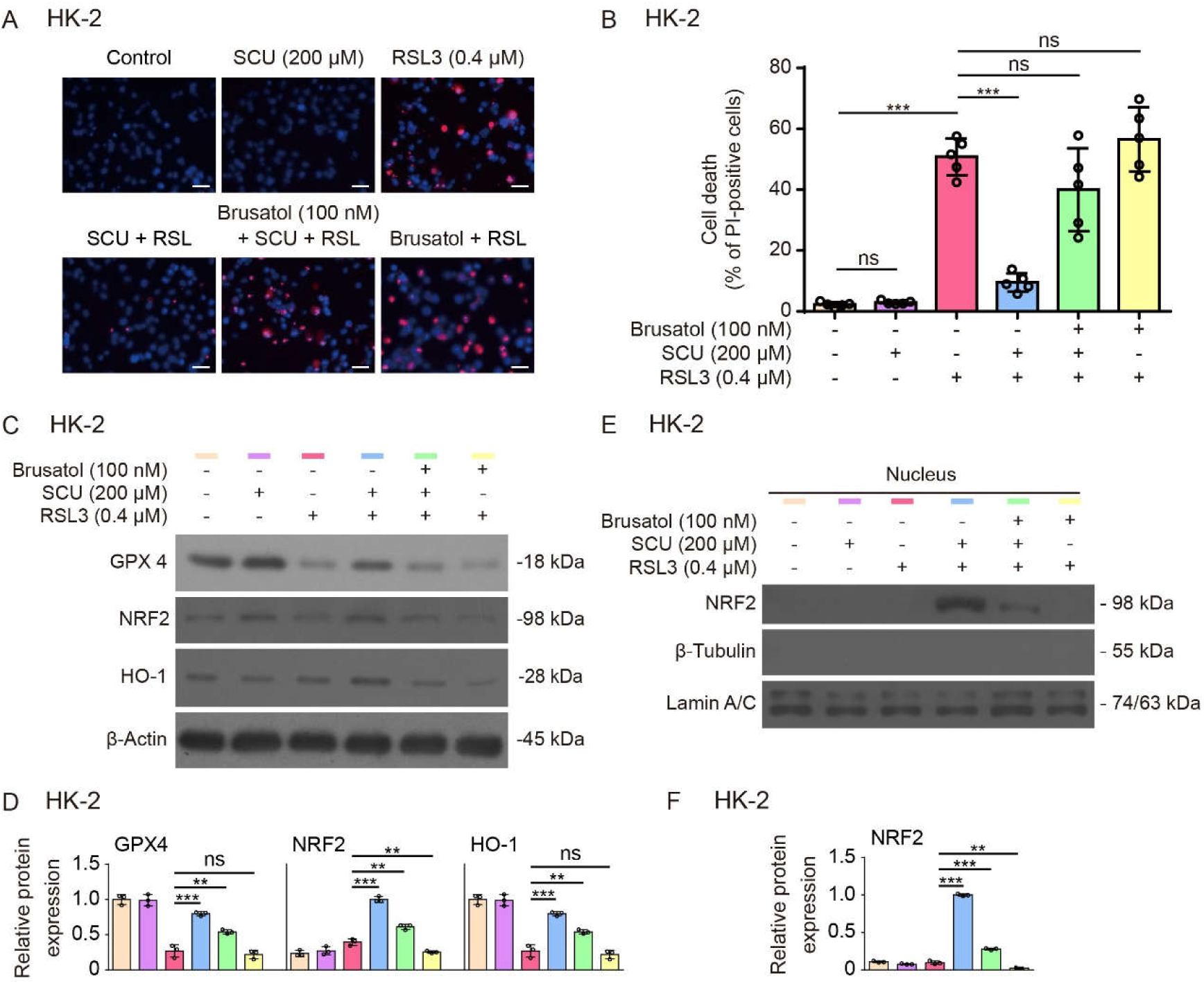
Scutellarin-mediated inhibition of ferroptosis is dependent on Nrf2 signaling. (A, B) HK-2 cells were pretreated with SCU alone or in combination with brusatol for 1 h, followed by treatment with RSL3 for 5 h. Cell death was assessed by PI and Hoechst 33342 staining, and fluorescence images were captured using fluorescence microscopy (A). Scale bars, 50 µm. Quantitative analysis of percentages of cell death (B). Data are presented as mean ± SD (*n* = 5). (C, D) Western blotting analysis of Nrf2, HO-1 and GPX4 levels with or without brusatol (C) and quantitative analysis of indicated proteins (D). (E, F) Immunoblot analysis of nuclear NRF2 protein with or without brusatol (E) and quantitative analysis of NRF2 protein levels relative to lamin A/C (F). Lamin A/C were used as a loading control for the nuclear fraction. β-Tubulin was undetectable in the isolated nuclear fraction. Data are presented as mean ± SD (*n* = 3). \*\**P* < 0.01; \*\*\**P* < 0.001; ns, not significant.

### Scutellarin administration mitigates folic acid-induced acute kidney injury

As ferroptosis is critical in renal injury in a mouse model of FA-induced AKI [33], we next assessed the effect of scutellarin on ferroptosis and renal injury in this mouse model. Acute renal injury was induced by a single dose of nephrotoxic FA intraperitoneally and scutellarin was orally administered thrice as indicated in **Figure 8A**. Histochemical analysis of kidney sections showed that FA induced marked pathological damage in the kidney, as revealed by pronounced vacuolization and increased Bowman’s space, which was ameliorated by scutellarin administration (**Figure 8B**). This was further supported by serum biochemical analyses, showing that scutellarin treatment significantly decreased serum levels of BUN and creatinine in mice challenged with FA (**Figure 8C, D**). These results together indicate that scutellarin can protect mice from FA-induced AKI.

**Figure 8.**
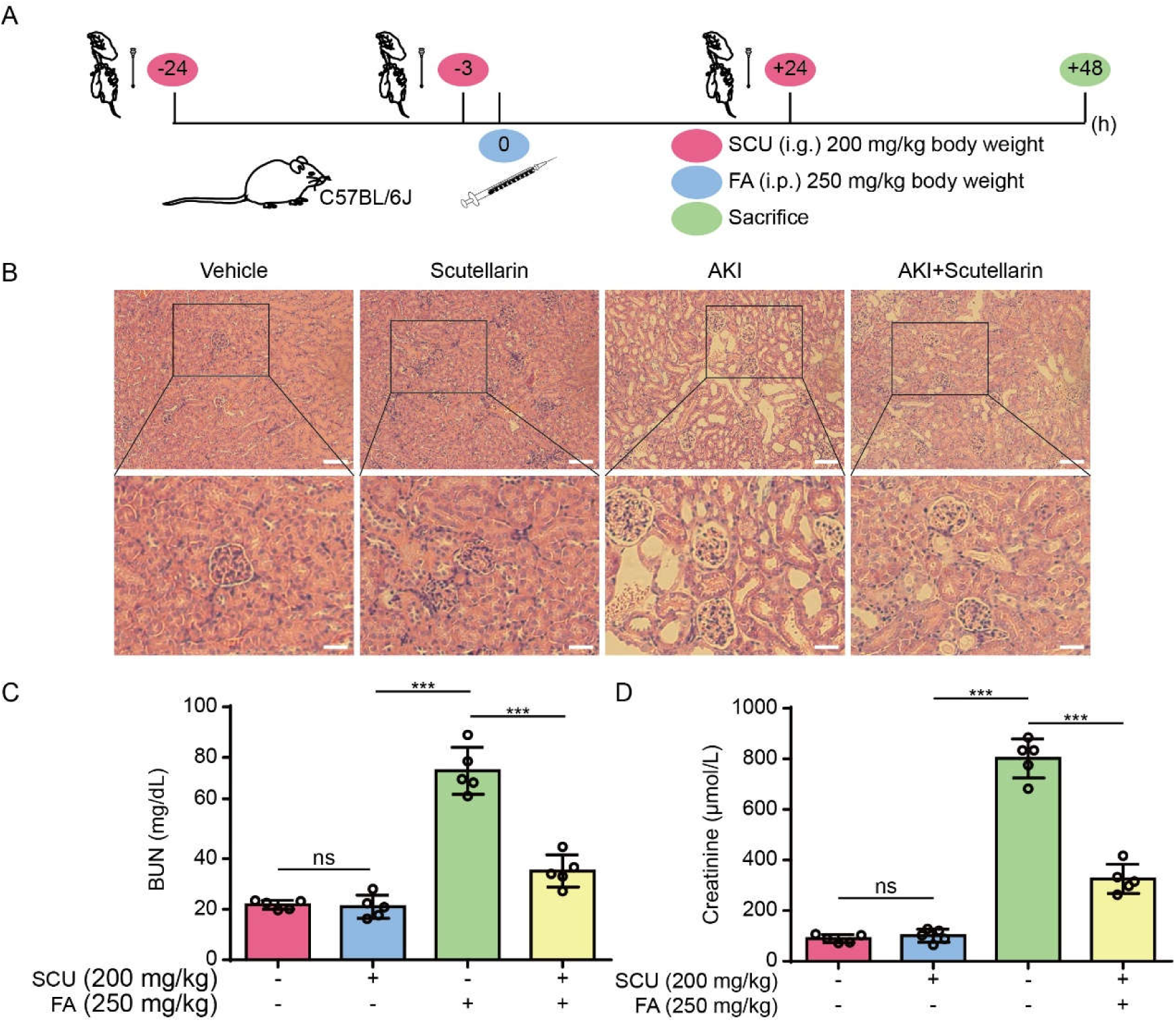
Folic acid-induced acute kidney injury is ameliorated by scutellarin. (A) Schematic depicting SCU administration timeline for the folic acid (FA)-induced mouse model of AKI. (B) Histochemical analysis (H&E staining) of kidney tissue sections from the AKI model. Scale bars, 50 µm (20 µm for insets). (C, D) Analysis of creatinine and blood urea nitrogen (BUN) levels. Data are presented as mean ± SD (*n* = 5). \*\**P* < 0.01; \*\*\**P* < 0.001; ns, not significant; i.p., intraperitoneal; i.g., intragastrical.

### Mitigation of acute kidney injury by scutellarin is associated with decreased ferroptosis signaling

To further assess an association between FA-induced AKI and ferroptosis, we analyzed the levels of GPX4, Nrf2, HO-1, and 4-HNE, the latter of which is a product of lipid peroxidation and is commonly used as a surrogate marker of ferroptosis [11, 34]. Western blotting showed that FA significantly decreased the levels of GPX4 in the kidney and markedly increased the levels of proteins modified by 4-HNE (**Figure 9A, B**), indicating lipid peroxidation and ferroptotic cell death in the kidney. In contrast, scutellarin administration increased GPX4 but decreased 4-HNE levels, indicating diminished lipid peroxidation and ferroptosis. Furthermore, Nrf2 and HO-1 levels were increased by scutellarin in the kidney of mice treated with FA (**Figure 9A, B**). We further assessed ferroptosis in cellular levels by detecting 4-HNE with immunofluorescence staining. Fluorescence microscopy showed that 4-HNE was pronouncedly increased in renal macrophages (MAC-2-positive) and other cells (**Figure 9C, D**), which was abrogated by scutellarin co-administration. Together, these results suggest that scutellarin mitigates FA-induced renal injury by inhibiting lipid peroxidation and ferroptosis in the kidney of mice, which is likely mediated by regulating the Nrf2 signaling.

**Figure 9.**
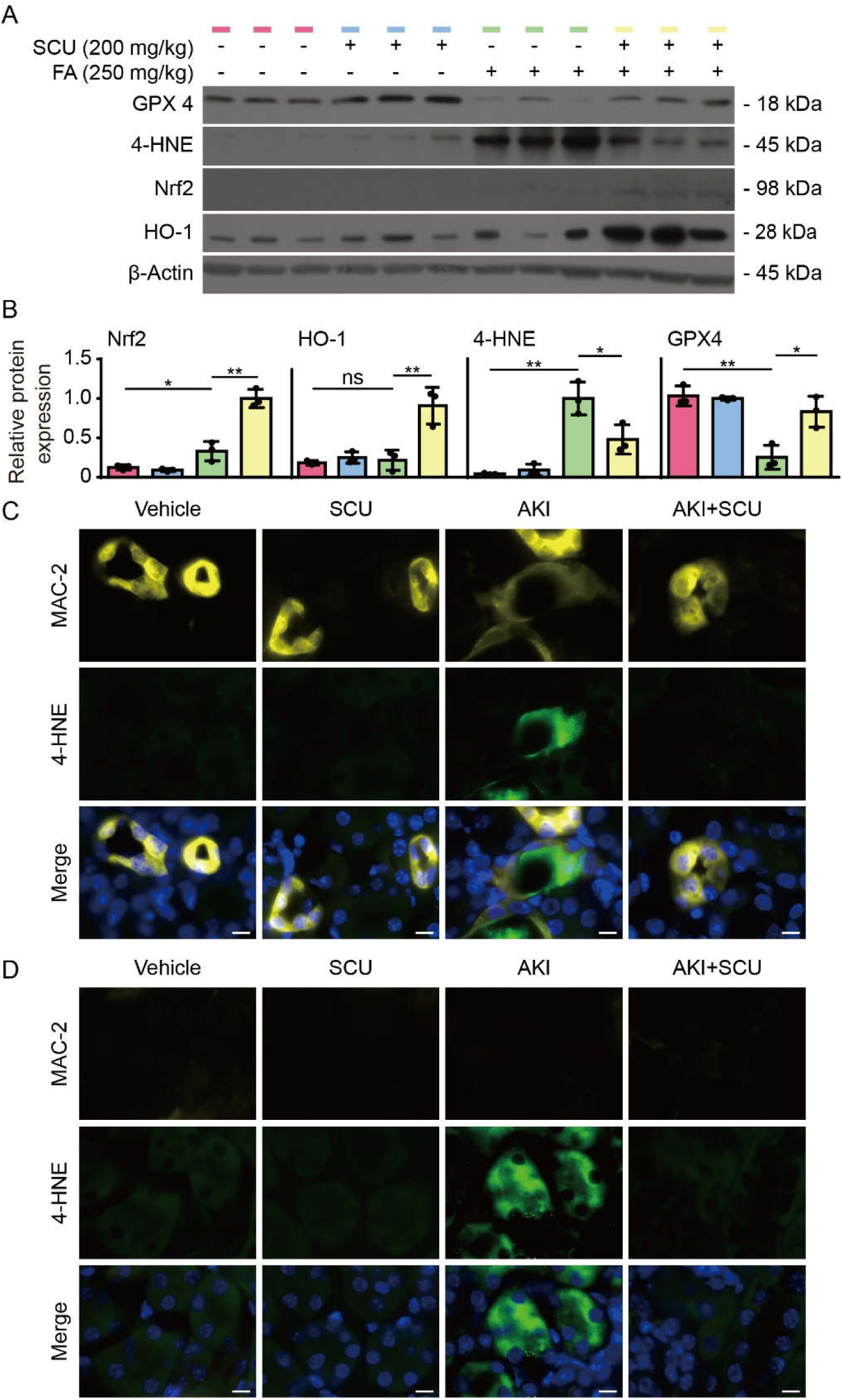
Mitigation of folic acid-induced acute kidney injury is associated with decreased levels of ferroptotic hallmarks in the kidney. C57BL/6J mice were treated as shown in Figure 7. (A) Western blotting was used to analyzed the expression of GPX4, 4-HNE, Nrf2, and HO-1 in the kidneys of 3 mice per group. β-Actin was used as a loading control. (B) Quantitative analysis of the expression levels of related proteins in (A). Data are presented as mean ± SD (*n* = 3). \**P* < 0.05; \*\**P* < 0.01; \*\*\**P* < 0.001. (C, D) Fluorescence microscopy show 4-HNE in macrophages (MAC-2 positive) and other cells (MAC-2-negative) in the kidney. Kidney tissues were fixed with 4% paraformaldehyde and frozen sections were prepared. After antigen retrieval, tissue sections were stained with specific primary antibodies and corresponding fluorescent secondary antibodies. Immunofluorescence images were captured using a fluorescence microscope. Scale bar, 10 µm.

## Discussion

Ferroptosis is a form of oxidative cell death that is caused by iron-dependent lipid peroxidation and can be induced by blocking cellular antioxidant systems such as the GSH/GPX4 axis and has been implicated in various pathological conditions including acute kidney injury [4, 6, 14]. In this study, we explored the effects of scutellarin on ferroptosis of human HK-2 cells and mouse primary BMDMs treated with RSL3 (a GPX4 inhibitor) and erastin (a system X_c_^−^ inhibitor), respectively. Our results showed that the flavonoid scutellarin was able to inhibit ferroptosis in these cells treated either with RSL3 or erastin in vitro and mitigate FA-induced renal injury in a mouse model of AKI. Our study highlights scutellarin as an inhibitor of ferroptosis and thus having potential applications for the treatment of ferroptosis-related renal injury.

Scutellarin is a flavonoid which is the major ingredient of breviscapine used for the treatment of cerebrovascular and cardiovascular diseases [19]. Previous studies indicate that scutellarin may exert its therapeutic effects on these diseases by its anti-inflammatory and antioxidative activities [18]. The therapeutic effects of scutellarin may also be mediated by its inhibitory activity on inflammasome-induced pyroptosis [21–23]. Our current study thus adds another layer of therapeutic mechanism for scutellarin by inhibiting the oxidative cell death—ferroptosis. Scutellarin inhibited ferroptosis in HK-2 cells and macrophages treated with either GPX4 inhibitor RSL3 or system X_c_^−^ inhibitor erastin. Interestingly, scutellarin could inhibit lipid peroxidation as revealed by C11-BODIPY 581/591 staining, suggesting that it might have acted on the upstream targets of ferroptotic pathways. Indeed, we found that scutellarin could counteract RSL3- and erastin-induced decrease in GPX4, the central antioxidant enzyme that can terminate the propagation of lipid peroxides [3]. Further, our data showed that scutellarin increased GPX4 levels by upregulating the Nrf2 signaling as blocking the activity of this transcription factor by Nrf2-specific inhibitor brusatol abrogated the effects of scutellarin. This is consistent with previous studies showing that scutellarin can upregulate Nrf2 activity [35, 36]. In further support of our results, the gene encoding GPX4 has been shown to be the target gene regulated by Nrf2 [12, 13]. In addition, the genes that encoding components of the system X_c_^−^ has also been regulated by Nrf2 [12, 13], but it is not known whether scutellarin has any effect on the system X_c_^−^. Together, our data reveal that scutellarin can counteract ferroptosis by upregulating the Nrf2 activity and thereby increasing the antioxidant capacity of the cell.

Mitochondria have critical roles in regulating ferroptosis [9, 30, 37]. It has been shown that mtROS contribute to ferroptosis by promoting lipid peroxides [29]. Consistent with this, we found that MMP was significantly reduced concomitant with increased production of mtROS during induction of ferroptosis, whereas scutellarin markedly suppressed mtROS production and prevented MMP reduction accompanied by inhibition of ferroptosis. Then how scutellarin protected mitochondrial dysfunction? One possibility is that scutellarin upregulated Nrf2 transcription factor, as mentioned above, thereby promoting the levels of the antioxidants, such as GPX1 and HO-1, of the cell to protect mitochondria. Of note, it has been found that DHODH operates in the mitochondrial inner membrane in parallel to mitochondrial GPX4 to inhibit ferroptosis by reducing ubiquinone to ubiquinol in certain cancer cells [9]; whether scutellarin can regulate this mitochondrial axis yet warrants further investigation. The second possibility is that scutellarin may attenuate mitochondrial damage through diminishing mitochondrial glucose oxidation by targeting the pyruvate dehydrogenase kinase (PDK)-pyruvate dehydrogenase complex (PDC) axis as reported previously [38]. A third possible mechanism is that scutellarin may act as a lipophilic radical-trapping antioxidant to terminate mitochondrial lipid peroxidation, which has been detected in mitochondria during ferroptosis [9, 10]. Yet, these possible actions of scutellarin need further clarification.

Our data show that scutellarin can inhibit ferroptosis by augmenting the antioxidant capacity of the cell by activating the Nrf2 signaling, thus being different from commonly used ferroptosis inhibitors such as Fer-1 and liproxstatin-1. It is known that Fer-1 inhibits ferroptosis by acting as a powerful radical scavenger and chain-breaking antioxidant, thus preventing lipid peroxidation and ferroptotic cell death [39]. Another potent ferroptosis inhibitor liproxstatin-1 acts to inhibit lipid peroxidation and to rescue GPX4 levels that can protect against lipid peroxidation [39]. Thus, scutellarin as a natural product represents an alternative candidate with different action mechanism for modulating ferroptosis-related disorders.

Consistent with in vitro results, our in vivo data showed that scutellarin was able to inhibit ferroptosis (as revealed by 4-HNE, a surrogate marker of ferroptosis) in macrophages and other renal cells of the kidney of FA-treated mice. Inhibition of ferroptosis was associated with reduced renal injury and increased expression of Nrf2, HO-1 and GPX4, suggesting that scutellarin might have exhibited its anti-ferroptotic activity by promoting the antioxidant capacity of cells in the kidney. Yet other mechanisms for scutellarin may also act in vivo in parallel to the upregulation of the Nrf2 signaling.

Although our in vivo results showed that oral administration of scutellarin was able to mitigate FA-induced AKI likely through inhibiting ferroptotic signaling, we currently lack direct evidence for the effective exposure of the kidney to scutellarin, which is a limitation of this study. Several previous studies have reported the pharmacokinetics and tissue distribution of scutellarin in mice, rats and humans following different routes of administration. For example, intravenously administered scutellarin (20 mg/kg body weight) could distribute into various organs including the kidney [40]. Another study showed that ⁓10% of the drug might be secreted through urine after a single oral dose of scutellarin (400 mg/kg body weight) in rats, suggesting a substantial part of the drug had been absorbed and eliminated by the kidney [41]. A clinical study further revealed that oral administered scutellarin (60 mg for average 64 kg body weight) can be absorbed into blood even though the plasma concentration of scutellarin was low [42]. These studies suggest that scutellarin can be absorbed and redistributed into the kidney even though its bioavailability is low. Thus, multiple administration of this drug seems necessary to achieve an effective concentration in targeted organs such as the kidney.

In conclusion, we show in this study that the flavonoid scutellarin can effectively inhibit ferroptosis in vitro in cultured cells and in vivo in the kidney of mice with FA-induced AKI. The Nrf2 signaling pathway has a critical role in mediating the anti-ferroptotic activity of scutellarin. Our data highlight that scutellarin can act as an inhibitor of ferroptosis to cope with ferroptosis-related diseases including acute renal injury.

