## Supplementary material for "Scutellarin inhibits ferroptosis by promoting cellular antioxidant capacity through regulating the Nrf2 signaling": Figure S

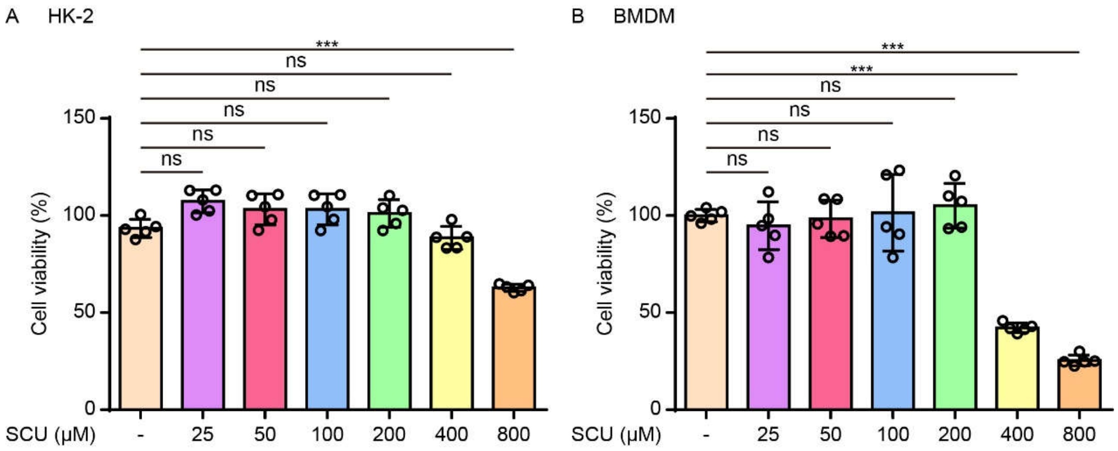


**Supplementary Figure S1. The cytotoxicity of scutellarin on HK-2 and bone marrow-derived macrophages (BMDMs).** Cells were treated with different concentrations of scutellarin (SCU) for 24 h, followed by addition of WST-1 reagent. The absorbance at 450 nm was measured using a microplate reader. The cell viability was presented as percentages of control for HK-2 cells (A) and BMDMs (B). Data are presented as mean ± SD ($n$ = 5). **$P < 0.01$; ***$P < 0.001$; ns, not significant.

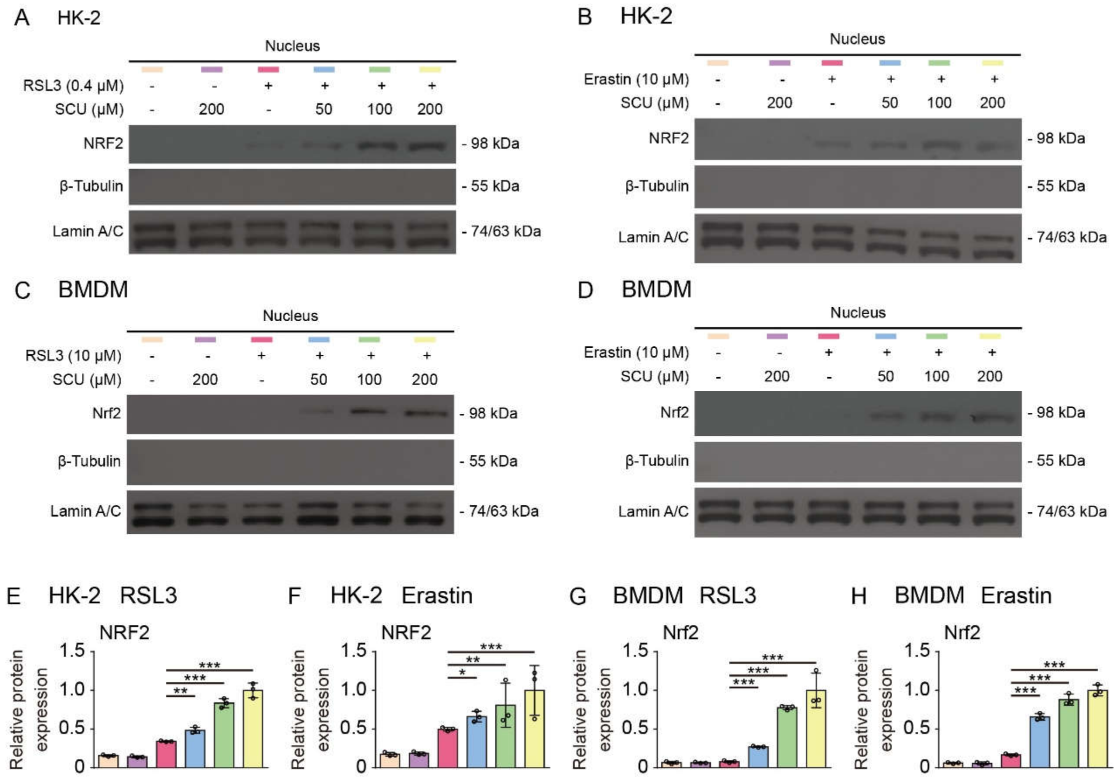


**Supplementary Figure S2. The effect of scutellarin on nuclear translocation of NRF2.** Cells were pretreated with different concentrations of scutellarin (SCU) for 1 hour, followed by treatment with RSL3 or erastin for 5 h (for HK-2) or 24 h (for BMDMs). (A-D) Western blot analysis of Nrf2 levels in the nuclear fraction. Lamin A/C were used as a loading control for the nuclear fraction. β-Tubulin was undetectable in the isolated nuclear fraction. (E-H) Quantitative analysis of protein levels of NRF2 relative to lamin A/C in (A), (B), (C), (D), respectively. Data are presented as mean ± SD ($n$ = 3). *$P < 0.05$; **$P < 0.01$; ***$P < 0.001$.
